# DNA Data Storage Architecture via Ligation of Dynamic DNA Bytes

**DOI:** 10.1101/2025.09.15.676197

**Authors:** Lijia Jia, Yue Shi, Jing Yang, Shangzhe Li, Wenjing Yang, Wei Li, Mancang Zhang, Quanshun Li, Yifei Zhang, Xiaolin Wang, Lin Li, Bo Duan, Dongbo Bu, Fei Chen, Haizhou Liu, Huaiyi Yang, Yongyong Shi, Di Liu

**Affiliations:** State Key Laboratory of Virology and Biosafety, Wuhan Institute of Virology, Chinese Academy of Sciences; Wuhan 430071, China; University of Chinese Academy of Sciences; Beijing 100049, China; Department of Microbial Physiological & Metabolic Engineering, State Key Laboratory of Microbial Diversity and Innovative Utilization, Institute of Microbiology, Chinese Academy of Sciences; Beijing 100101, China; Beijing Key Laboratory of Genetic Element Biosourcing & Intelligent Design for Biomanufacturing; Beijing 100101 China; United Research Center for Next Generation DNA Synthesis of SJTU Dynegene; Shanghai 201108, China; Bio-X Institutes, Shanghai Jiao Tong University; Shanghai 200030, China; Key Laboratory for Molecular Enzymology and Engineering of Ministry of Education, School of Life Sciences, Jilin University; Changchun 130012, China; Sanya Institute of Hunan University of Science and Technology; Sanya 572024, China; School of Life and Health Sciences, Hunan University of Science and Technology; Xiangtan 411201, China; State Key Laboratory of Pathogen and Biosecurity, Beijing Institute of Microbiology and Epidemiology; Beijing 100071, China; Institue of Computing Technology, Chinese Academy of Sciences; Beijing 100190, China; Western Institute of Computing Technology; Chongqing 401121, China; China National Center for Bioinformation; Beijing 100101, China; Beijing Institute of Genomics, Chinese Academy of Sciences; Beijing 100101, China; Institute of Neuroscience, Center for Excellence in Brain Science and Intelligence Technology, Chinese Academy of Sciences; Shanghai 200031, China

**Keywords:** DNA data storage, dynamic DNA bytes, CRUD operations, hierarchical access, error correction

## Abstract

The explosive growth of digital data is overwhelming conventional storage media, creating an urgent need for more efficient solutions. DNA offers immense potential for digital data storage, yet most systems remain static and archival. Here, we present a modular DNA storage architecture based on dynamic DNA bytes (DynaBytes)— pre-fabricated DNA segments that can be ligated into reconfigurable information units. Utilizing core, functional and control DynaBytes, we stored 210,776 bits (26,347 bytes) of digital information organized within a file-system, and demonstrated CRUD (Create-Read-Update-Delete)-like operations, hierarchical access and nanopore-based realtime retrieval. Robust data recovery was achieved under ∼100x error-prone sequencing using streamlined error correction and fuzzy decoding. By relying on *in vitro* ligation of standardized components, DynaBytes reduces cost, scales efficiently, and allows interactive, rewritable storage. These features advance DNA storage beyond passive archiving toward a reconfigurable framework, opening new possibilities for dynamic, practical and large-scale DNA-based data systems.

## Introduction

The exponential expansion of digital information—driven by AI, IoT and high-throughput data generation—has begun to approach the physical limits of conventional silicon-based storage in terms of capacity and sustainability [1–3]. DNA, nature’s native data carrier, has emerged as a promising alternative medium for digital information storage, offering ultrahigh information density, long-term stability and energy-efficient preservation [4–6]. Over the past decade, a range of proof-of-concept studies have demonstrated the feasibility of writing, preserving and reading digital information using synthetic DNA, laying a foundation for molecular-scale information systems [5, 7, 8].

Two main technical paradigms have shaped the field of DNA data storage. The first and most established approach relies on de novo synthesis of data-encoding strands using dense coding strategies [9–11]. These approaches achieve high information density and robust recovery through advances in sequence design and error correction [12, 13], making them well suited for archival applications. However, their write-once nature, high synthesis cost and lack of direct access to individual molecules remain key limitations for dynamic or large-scale use [6, 14]. In contrast, a modular strategy inspired by movable type has gained traction in recent years: instead of synthesizing new strands for each file, systems pre-fabricate a finite set of DNA fragments that can be combinatorially assembled to encode digital data [15–17]. This strategy reduces synthesis burden, allows for parallelized writing and offers greater flexibility for reconfiguration. Yet, current implementations typically support only simple data layouts and lack the dynamic capabilities needed for real-world applications.

Despite technical progress in encoding [18–20], error correction [21, 22] and decoding [23], DNA data storage systems still fall short of providing the interactivity, scalability and programmability required for functional molecular information systems [1]. Bridging this gap calls for reimagining DNA not only as a dense medium for static storage, but also for flexible information processing—supporting operations like search, editing and hierarchical data organization within a unified biochemical framework. Here, we present Dynamic DNA Byte (DynaByte), by which a modular DNA data system was designed to support flexible storage, efficient retrieval and structured file management. Advanced to the our previous prototype [15], DynaByte introduces a hierarchical and reusable set of DNA components. At its core are addressable storage units—address DynaBytes, content DynaBytes and error correction code (ECC) DynaBytes—which together support fundamental write and read operations. On top of this foundation, a set of functional DynaBytes were implemented to encode metadata and enable logic-based control: version DynaBytes allow file editing and versioning, filename DynaBytes support name-based file retrieval and carrier DynaBytes offer a linear plasmid backbone with directory labels for organizing files hierarchically. Combined with an optimized decoding pipeline, DynaByte enables key functionalities such as data manipulation, Oxford Nanopore Technology (ONT)-based data retrieval and hierarchical file access (Fig. 1a). Using this system, we exemplified storing 210,776 bits (26,347 bytes) digital information, demonstrating its potential as a scalable and interactive platform for DNA data storage.

**Fig. 1.**
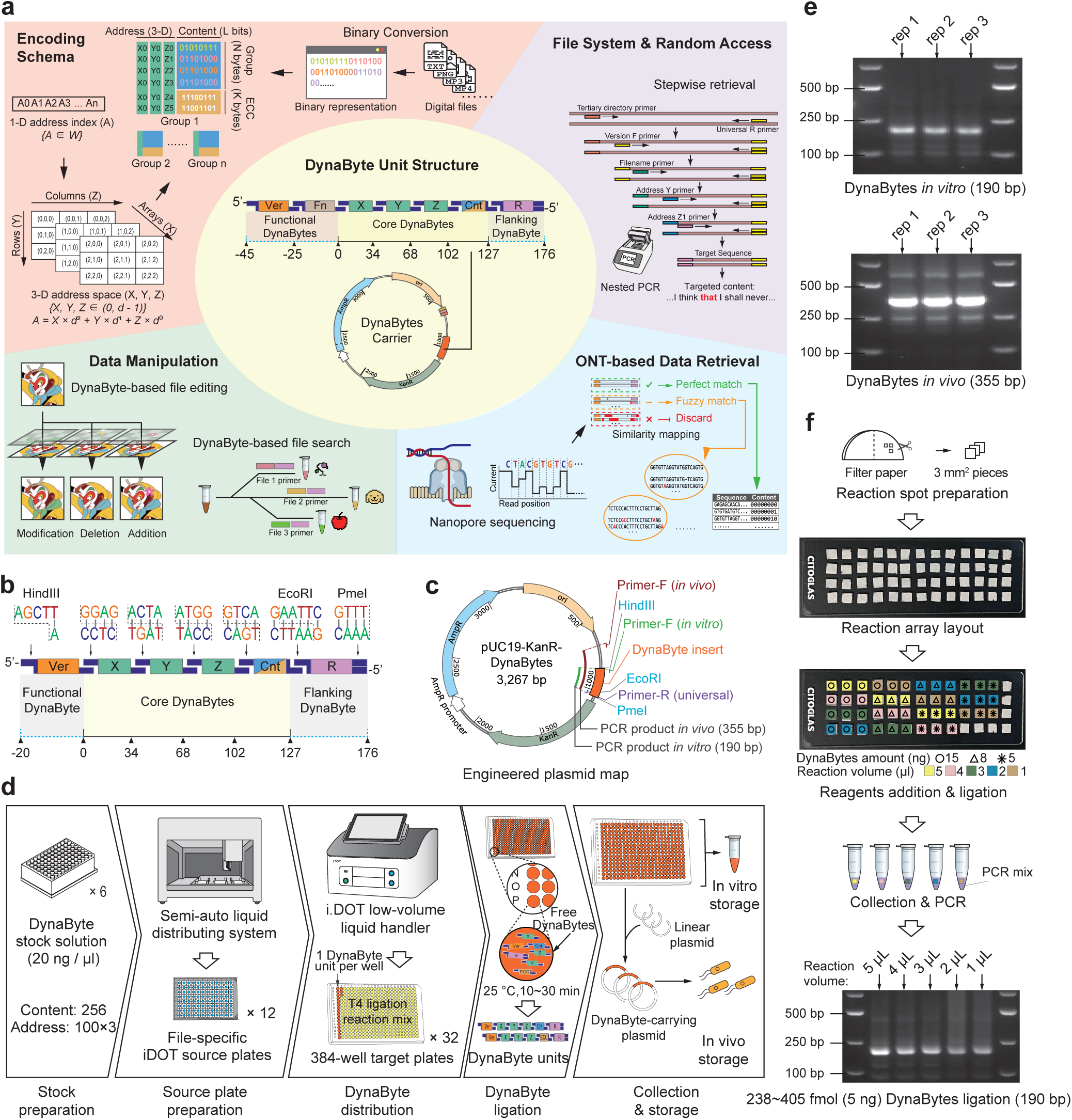
Schematics and Overall Design of the DynaByte Storage System. **a**, Overview of the DynaByte storage system. **b**, Structural design of a DynaByte unit, comprising version information, X-Y-Z addresses, content region and a right-flanking DynaByte. Linker sequences connecting adjacent DynaBytes are illustrated. **c**, Engineered plasmids used to carry DynaBytes. Primer pairs used for amplification and expected amplicon sizes are indicated. **d**, Workflow of high-throughput DynaByte storage. Parallelized processing is enabled by an automated liquid dispensing system (see Figure S2 for setup). **e**, 1% agarose gel electrophoresis results showing successful amplification of DynaByte units both *in vitro* and *in vivo*. **f**, Evaluation of ultra-miniaturized reaction systems for DynaByte storage. Storage feasibility was tested under varying DNA input amounts (ng) and enzyme reaction volumes (μL).

## Results

### Design and Implementation of the DynaByte Storage Architecture

Building upon “movable type” prototype [15], we present a scalable, modular and manipulable architecture termed DynaByte storage (Fig. 1a). Each DynaByte is a short, double-stranded, pre-synthesized DNA fragment (20–35 bp), designed to avoid homopolymers, high GC content and strong secondary structures (See Methods). Multiple DynaBytes can be concatenated via programmable enzymatic ligation using sticky ends, forming combinatorial storage units namely DynaByte Units encoding entire files or logical data blocks (Fig. 1b). The core components of each DynaByte Unit are three address DynaBytes (address-bytes) (X, Y, Z) and one content DynaByte (content-byte), collectively referred to as core DynaBytes. These fulfill the basic functionality required to store and retrieve digital data. Additional modular elements include a flanking DynaByte (flanking-byte, R), used for amplification and functional DynaBytes (functional-bytes) encoding logical operations such as version control (Fig. 1b). We designed and synthesized 256 distinct content-bytes and 300 address-bytes forming a hierarchical X-Y-Z indexing structure (Fig. S1a, Table S1). This design supports megabyte-scale (100³ bytes) data storage. Address-bytes adopt a two-part design, with shared prefixes enabling group-wise organization and distinguishable suffixes ensuring specificity (Fig. S1b).

The DynaByte Unit is compatible with both *in vitro* and *in vivo* storage. It can be embedded into engineered plasmids as circular DNA constructs (Fig. 1c), providing enhanced structural stability and enabling bacterial transformation. To support high-throughput synthesis and writing, we constructed a semi-automated DynaByte storage platform combining a custom-built stock-handling workstation with a non-contact liquid dispenser (I.DOT, Dispendix GmbH), allowing parallelized ligation reactions in nanoliter-to-microliter volumes (Fig. 1d, Fig. S2). Within the modular automation platform, DynaByte components at different levels were ligated in separate wells to form circular constructs, facilitating downstream cold storage or bacteria-based archival. We validated DynaByte assembly under standard (10 μL) and ultra-miniaturized (1 μL) reaction conditions. Gel electrophoresis confirmed successful assembly of the DynaByte Unit with clear target bands observed from both *in vitro* and *in vivo* samples (Fig. 1e). To further optimize reaction scalability, we tested a glass-slide “drop-on-paper” setup and determined that as little as 238∼405 fmol of each DynaByte was sufficient to complete ligation in 1 μL total volume (Fig. 1f), supporting the feasibility of large-scale, low-input DNA data encoding.

### Robust Data Encoding with DynaByte Outer Error Correction Code

Instead of the commonly-used two-dimensional Reed–Solomon (RS) error correcting strategy [11, 21], DynaByte storage applied a one-dimensional RS based outer code, termed the DynaByte Outer Code. This scheme introduces a new class of redundancy symbols called ECC DynaBytes (ECC-bytes), which are derived from RS symbols and stored alongside original content-bytes to enable error correction (Fig. S3a, Methods). This one-dimensional, content-oriented design simplifies implementation while retaining robustness. Simulations-based optimization determined the RS group size (*k*) and the number of RS symbols (*n*) per group. Two test files—a 1.18 KB text and a 2.62 KB image (converted to 3.51 KB base64 format)—were encoded using the DynaByte system (Fig. 2a). We selected *k = 4* and *n = 2* (50% redundancy) to balance recovery performance and overhead under varying dropout rates (Fig. S3b). After byte-encoding, grouping and ECC-bytes embedding, the files expanded to 1,824 and 5,388 bytes, respectively (Fig. 2b; Table S2–3). Content-byte usage showed uniformity for text but structured bias for the image, suggesting potential for format- or content-aware optimization (Fig. 2c). Address-byte prefixes displayed low complexity, reflecting group-wise organization (Fig. 2d). Most reads retained full-length core DynaBytes (127 bp, Fig. 2e) and 100× subsampling resulted in only a small number of missing units—3 for the text file and 8 for the image (Fig. 2f). Additionally, while most index-addressed pairings correctly matched their target content-bytes, a few false-positive pairings (“spurious content-bytes”) displayed stronger signal than the true targets, leading to potential misidentification (Fig. 2g). Despite observed errors—base-level errors (leading to imperfect unit lengths), unit dropouts (coverage loss) and spurious content-byte signals (false positives mismatched with index addresses)—the combination of DynaByte encoding and ECC-byte redundancy enabled full data recovery. By developing a DynaByte decoding algorithm and simulating varying sequencing depths, we found that strict exact decoding achieved 100% data recovery at 200×, while fuzzy decoding succeeded at just 100× (Fig. 2h, Fig. S3c, Table S4). These results underscore the robust error-handling capacity of the ECC-augmented DynaByte system under realistic sequencing conditions.

**Fig. 2.**
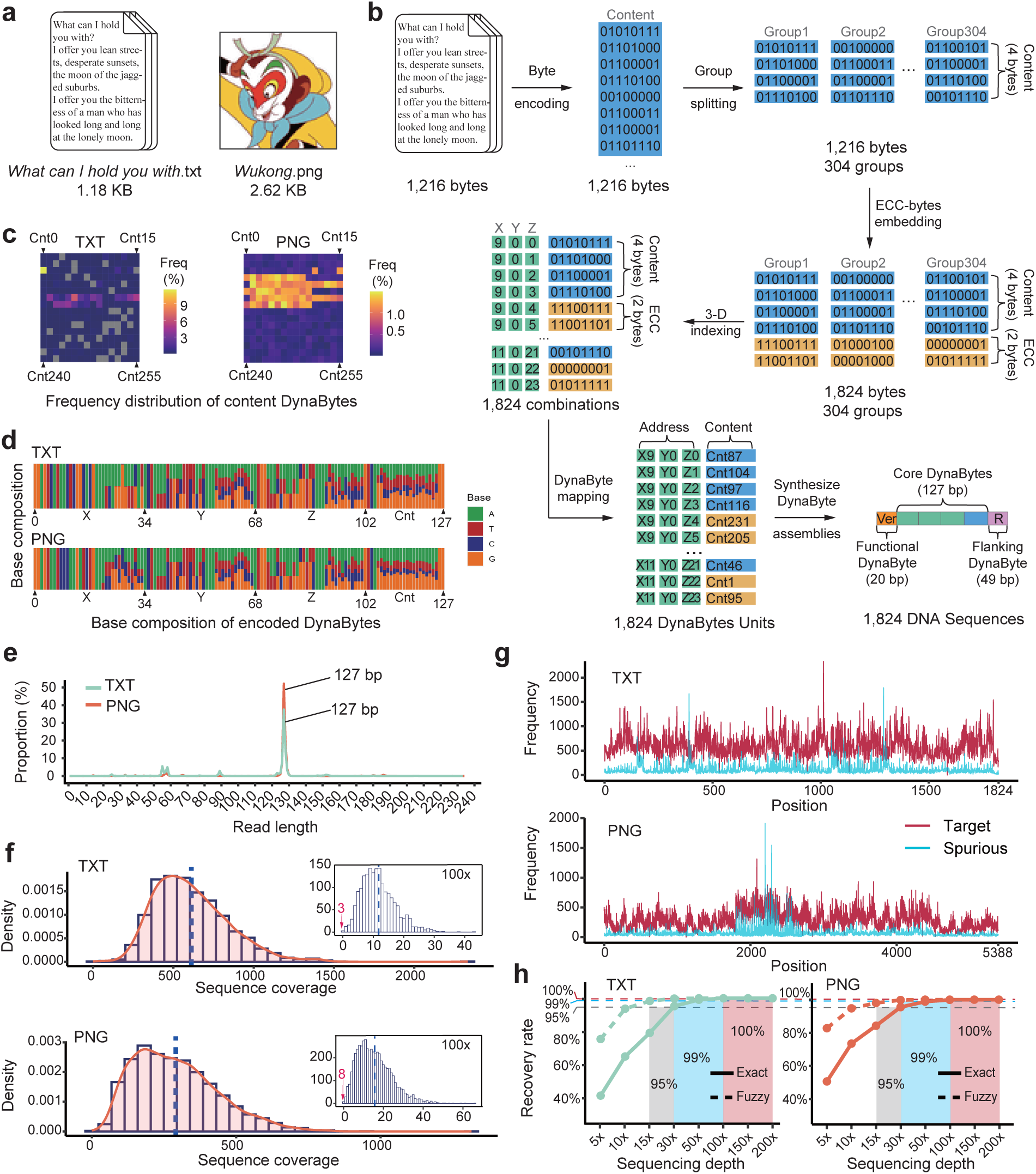
Basic Demonstration and Performance Assessment of DynaByte Storage and Data Retrieval with ECC-bytes. **a**, Example files (TXT and PNG) used for demonstration of DynaByte-based storage. **b**, Schematic of content element grouping and Error correction code DynaByte (ECC-byte) encoding. The original bitstream is first byte-encoded, meaning it is divided into content elements of fixed length, with each element consisting of m = 8 bits (i.e., one byte). This yields a total of L = (length of bitstream)⁄m elements. The elements are then grouped into blocks of k elements each, resulting in K = ⌈L⁄k⌉ groups. The final group, if shorter than k, retains its actual length and is still processed for error correction. Each group, with actual length k, is encoded using a (ki+n, ki) Reed-Solomon (RS) code to generate n ECC-bytes for error correction. This design enables up to ⌊n⁄2⌋ symbol-level error correction per group. In this example, *m* = 8, *L*=1216, *k* = 4, *K*=304, *n* = 2. Each group is thus encoded into 6 bytes, including 2 ECC-bytes. **c**, Heatmaps of content DynaByte usage across the two example files, showing the frequency of each DynaByte in the encoded data. **d**, Nucleotide composition of the DynaByte-encoded sequences for the two files. **e**, Length distribution of sequencing reads after trimming off the universal flanking DynaBytes (Ver and R). **f**, Coverage of encoded sequences in the raw sequencing data and after 100× subsampling. The number of undetected sequences is labeled in red. **g**, Signal-to-noise ratio of sequencing results. Each index address corresponds to a content DynaByte. True positives (“Target”, magenta) match the expected encoded sequences; false positives (“Spurious”, cyan) represent incorrect content DynaBytes. **h**, Simulated data recovery performance at varying sequencing depths. Two decoding strategies are compared: Exact decoding (strict sequence matching) and Fuzzy decoding (tolerant to minor errors). The recovery rates of 95%, 99% and 100% are marked by dashed lines, with corresponding depth intervals highlighted in color.

### Programmable Data Manipulation by using Functional DynaBytes

Unlike digital storage systems where editing a file is direct and efficient, DNA storage typically lacks native mechanisms for content modification—editing usually requires rewriting the entire dataset. To address this limitation, we developed a programmable manipulation strategy in the DynaByte system by introducing functional DynaBytes (functional-bytes), enabling database-like operations such as modification, deletion, addition and targeted retrieval.

To demonstrate in-storage data manipulation, we designed version DynaBytes (version-bytes) to support versioning and file-layer overlay. Using *Wukong.png* as a test file, we generated three editing layers: scarf color modification, hairband deletion and flower addition (Fig. 3a). Each layer was encoded independently and labeled with a version-byte indicating the specific edit, then pooled with the original file in a common DynaByte library (Fig. 3b, Table S3). This modular strategy reduced storage burden by 61.2% compared to full-file rewrites (Fig. 3c). During retrieval, version-specific or multiplex PCR using multiplexed keys (primers) allowed users to selectively amplify and reconstruct the desired version. Gel electrophoresis confirmed that the PCR products exhibited main bands at the expected lengths (Fig. 3d, Fig. S4a). Subsequent NGS sequencing reveled that the sequence frequency distribution among different layers is quite similar to their encoded proportions, supporting the high fidelity and effectiveness of the version-byte strategy (Fig. 3e). Decoding these sequences yielded correctly reconstructed versions of the file, each reflecting the intended modifications—scarf color change, hairband deletion and flower addition—thereby validating the system’s capacity for programmable edit operations (Fig. S4b, Fig. 3f). Based on this, We conducted *in silico* down-sampling experiments to evaluate the recovery accuracy of each version under varying sequencing depths. Downsampling simulations further demonstrated that 99% recovery was achievable at 100× depth across all layers, and as low as 5× for some (Fig. 3g, Fig. S4c).

**Fig. 3.**
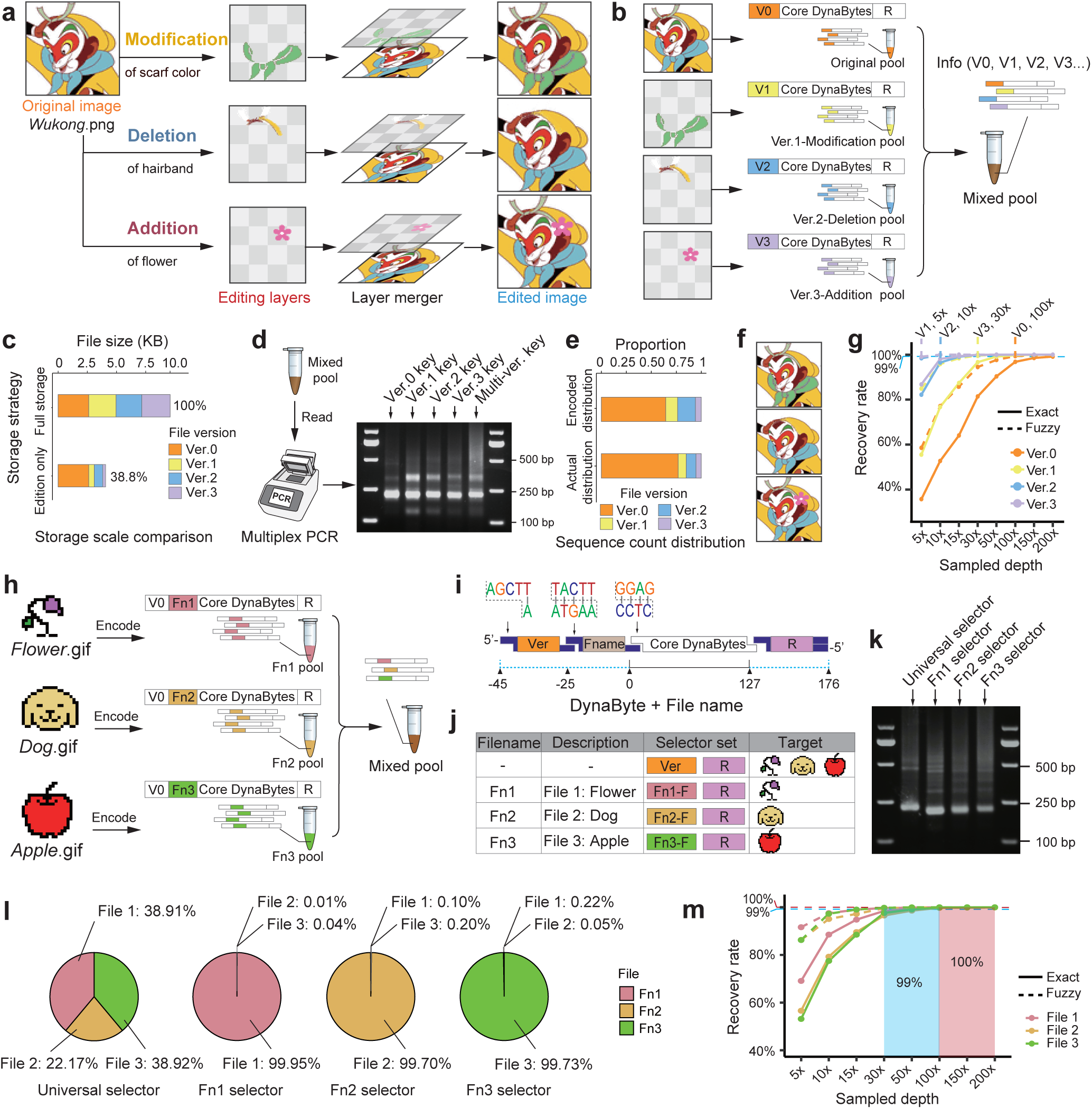
DynaByte Storage for Data Manipulation. **a**, Image file editing demonstrated via layered storage. **b**, Design of DynaByte storage structure enabling image addition, deletion and modification. Different version DynaBytes (version-byte) are used to label the version of each stored layer, which are then co-stored with the original image. **c**, Comparison of required storage size between this method and conventional approaches. **d**, Retrieval of version-specific content via multiplex PCR using key primers targeting version-bytes. **e**, Theoretical vs. observed proportion of sequencing reads corresponding to each version label in the encoded data. **f**, Decoded and recovered files corresponding to three types of edits: scarf color modification, hairband deletion and flower addition. **g**, Mean data recovery rates of different versioned content under varying sequencing depths (10 random replicates per depth). The 99% recovery threshold is shown as a dashed line, with the sequencing depth required to reach 99% recovery for each version specifically annotated. **h**, Design principle of file search using filename-specific DynaBytes. Multiple files are labeled with distinct filename DynaBytes (fname-bytes) and co-stored in a mixed pool. **i**, Structure of the expanded DynaByte unit after integration of fname-bytes. **j**, Comparison of content retrieval using either non-specific selectors or file-specific selectors targeting fname-bytes. **k**, Agarose gel electrophoresis results of amplification products from file selection. **l**, Distribution of recovered sequences across files based on sequencing data for each search group. **m**, Mean decoding rates of file-specific content under varying sequencing depths (10 random replicates per depth).

For file-level addressing, we implemented filename DynaBytes (fname-bytes), enabling targeted retrieval of individual files within a mixed pool. We encoded three GIF files, each labeled with a unique fname-byte positioned between the version-byte and the core-bytes, with standardized linkers ensuring consistent ligation order (Fig. 3h–i, Table S5). Selective amplification was achieved using selector primers corresponding to either fname-bytes (for single-file access) or version-bytes (for bulk retrieval) (Fig. 3j). Clear main bands observed in electrophoresis (Fig. 3k, Fig. S4d) confirmed that fname-bytes did not impair ligation efficiency. NGS analysis showed that over 99% of sequences retrieved by fname-byte selectors matched the correct target (Fig. 3l). On average, 99% recovery was achieved at 30× depth, and 100% at 100× for all files (Fig. 3m, Table S7).

Together, these functional-bytes provide the DynaByte system with data editing and querying capability, enabling CRUD (Create-Read-Update-Delete)-like operations and supporting scalable, selective and efficient DNA-based information management.

### DynaByte Decoding by Nanopore Sequencing

For data retrieval, we employed ONT sequencing to successfully decode information stored in DynaByte storage. As a real-time, portable Sequencing platform with inherently higher error rates than NGS, ONT challenges conventional encoding methods [24–26]. Base-bit mapping strategies are largely incompatible with ONT due to bit-level ambiguity caused by high sequencing noise [27, 28]. In contrast, DynaByte’s fixed structure and restricted codebook offer inherent error tolerance, making it well-suited for noisy sequencing environments. Using the file *what can I hold you with.txt* as an example, we implemented a decoding workflow based on fuzzy matching: after extracting DynaByte-structured reads, Levenshtein distance (LD) was used to match reads to the closest codebook entries (Fig. 4a, Fig. S3c). Compared to Illumina NGS, standard ONT showed higher LDs, however, the use of the high-accuracy sup basecalling model (ONT-sup) greatly reduced LDs, approaching NGS-level performance (Fig. 4b, Fig. S5a–d). Error quantification revealed that ONT reads had the highest average number of errors per read (8.05; Fig. 4c–d), with hotspots concentrated in the address-bytes X and first part of Y, suggesting that low-complexity sequences are major contributors to recognition errors (Fig. S5e). ONT-sup outperformed the standard ONT model in both error count and type but still fell short of NGS. We further assessed data recovery rates under varying sequencing depths (10 replicates per depth). Exact matching proved ineffective on ONT (only 19.58% recovery at 200×), whereas fuzzy decoding achieved 99.83% recovery at the same depth and full recovery at higher coverages (Fig. 4e, Table S7). At ≥100× depth, ONT-sup with fuzzy decoding matched NGS in recovery performance, with complete restoration of the stored content. These findings highlight DynaByte’s high platform compatibility and error resilience, particularly its suitability for fast, portable and noisy readout environments like ONT. The ONT-sup model, in particular, enhances decoding reliability and underpins DynaByte’s potential for real-world deployment on long-read sequencing platforms.

**Fig. 4.**
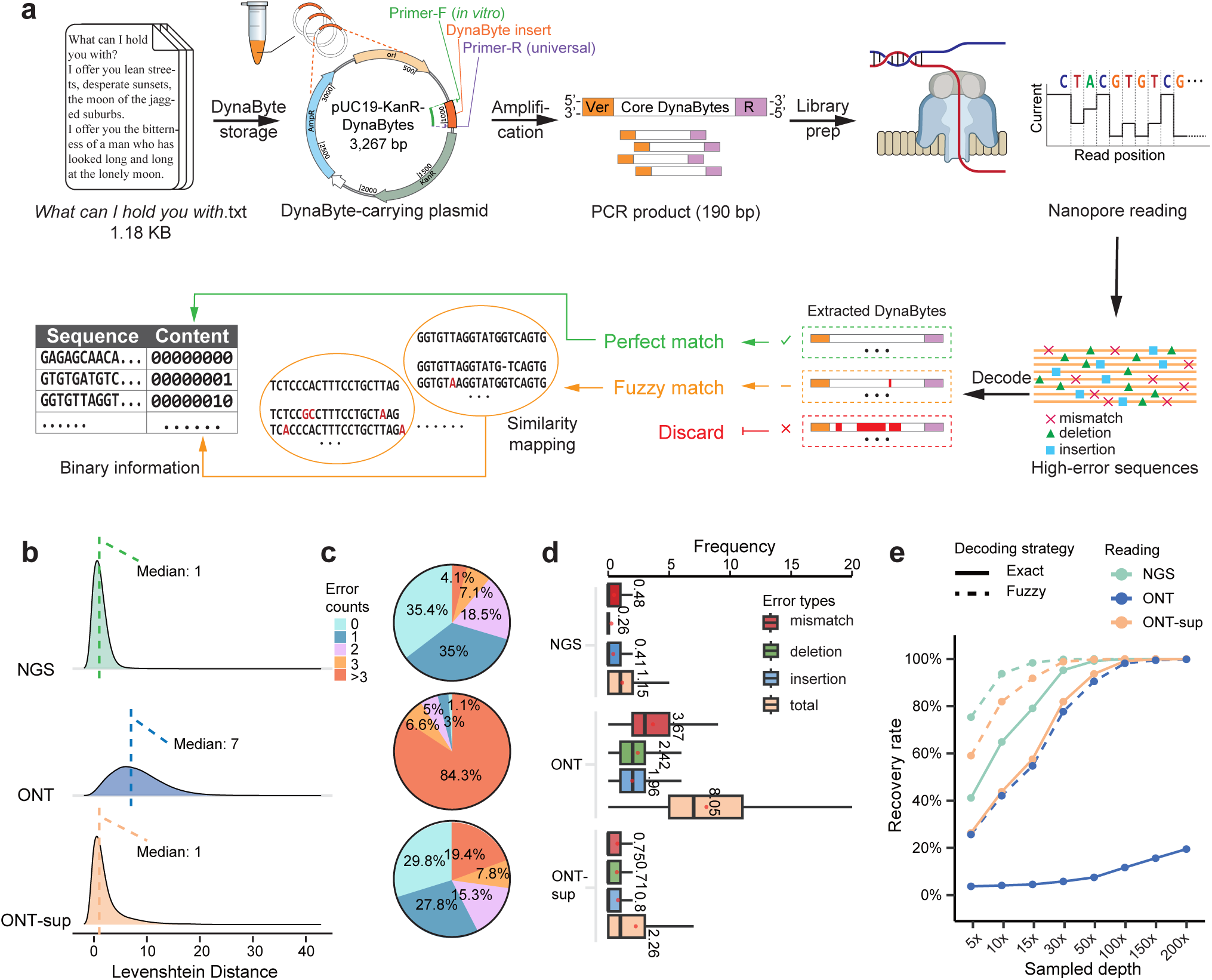
Data Retrieval from DynaByte Storage Using ONT Sequencing. **a**, Workflow for retrieving data from DynaByte storage using Oxford Nanopore Technologies (ONT) sequencing, along with the decoding strategy. Fuzzy matching calculates the Levenshtein distance between each sequencing read and the DynaByte codebook via DynaByte mapping, returning the closest match. **b**, Distribution of Levenshtein distances between sequencing reads and reference sequences for different methods: Illumina NGS, ONT and ONT with the high-accuracy sup basecalling model (ONT-sup). Medians are indicated on the plots. **c**, Distribution of sequencing errors per read across different retrieval methods. Each segment shows the proportion of reads with exactly x errors (x = 0, 1, 2, …). **d**, Comparison of error rates for the same file across different reading methods. Box colors indicate error types, and red dots represent the mean error rate for each type. **e**, Mean recovery rates of stored data at varying subsampling depths (10 random replicates per depth), comparing ONT and NGS-based methods.

### Hierarchical Access and Retrieval within the Prototype DynaByte File System

To enable hierarchical file organization and addressable data access within DNA storage, we extended the DynaByte system by introducing a three-level directory tagging scheme (3DTag), resulting in the construction of DynaByte Pro (Fig. 5a). The 3DTag comprises three 24-bp sequences: a primary directory (D1) corresponding to the native plasmid backbone, and two engineered sequences (D2 and D3) introduced via genetic modification. D1 serves as the physical root address, while D2 and D3 provide logical hierarchy and flexible classification (Fig. 5a, 5b). To validate the system’s capabilities in structured storage and hierarchical retrieval, we organized 10 digital files into a directory tree that mimics a conventional computer file system, encompassing directories, version control and file/address blocks (Fig. 5b, Table S6). Specific selectors (primers) targeting hierarchical metadata were designed, enabling a simulated set of hierarchical retrieval commands. Using semi-nested PCR, DynaByte Pro achieved targeted amplification of specific content blocks by directory, version, or address without sequencing the entire library (Fig. 5b, Fig. S6a). Experiments confirmed that this method selectively retrieved target content from mixed libraries at multiple retrieval granularities (Fig. 5c, Fig. S6b). Sequencing the PCR products revealed that, after trimming selectors and tags, the recovered sequence lengths closely matched the designed payloads (Fig. 5d, Table S8), validating the system’s accuracy and predictability. This method enables file- or subfile-specific reading without reliance on full-library sequencing, making it particularly suited for scalable, high-throughput DNA data access.

**Fig. 5.**
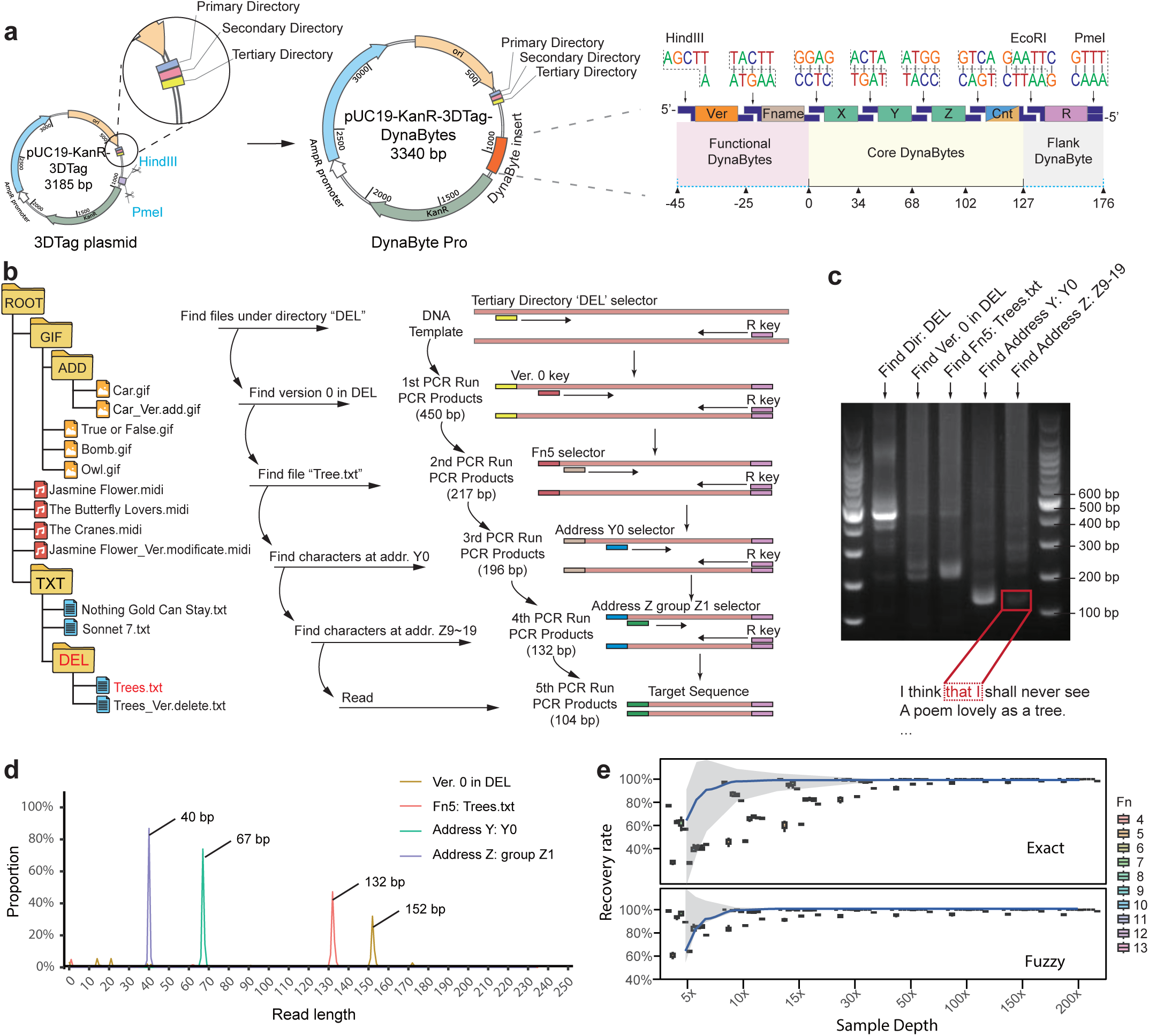
Hierarchical File Organization and Random Access enabled by the DynaByte Pro Storage System. **a**, Construction of a functional DynaByte storage system termed DynaByte Pro, which extends the core DynaByte framework by incorporating 3-level directory tags (3DTag) into a plasmid vector. **b**, Directory tree structure designed for DynaByte Pro and schematic illustration of stepwise content retrieval using semi-nested PCR guided by directory-, version-, filename- or address-specific selectors. **c**, Agarose gel electrophoresis results of semi-nested PCR products. **d**, Length distribution of decoded sequences after removal of flanking-bytes and selectors (retrieval primers) from the PCR products. **e**, Boxplot of decoding rates for 10 files stored in DynaByte Pro under varying subsampling depths (10 random replicates per depth). Gray shading indicates the interquartile range (IQR), and the line represents the median.

Further evaluation demonstrated robust decoding performance across a range of sequencing depths. We conducted 10 independent random decoding trials for each file at depths from 5× to 200×, assessing recovery rates under both fuzzy and exact decoding strategies (Fig. 5e, Table S7). Based on the median recovery rates across all files, fuzzy decoding achieved full recovery at depths as low as 30×, whereas exact decoding required ≥100× depth for full recovery. Although individual file performance varied at low depths, overall trends highlighted the superior robustness and error tolerance of the fuzzy strategy, particularly under low-coverage conditions. This feature is critical for cost-effective, scalable DNA storage applications.

Together, these results demonstrate that DynaByte Pro realizes a biochemical prototype of a digital file system, incorporating hierarchical organization, modular design and addressable random access. Beyond simple data storage, DynaByte units now support operations such as integrity checking, file segmentation and logical addressing, establishing a foundation for future high-performance DNA file systems.

## Discussion

Here, we present a modular, operable and scalable prototype of a molecular file system based on pre-designed double-stranded, ligation-ready DynaBytes. The system supports both database-like and file-level operations—including additions, deletions, modifications and hierarchical access—marking a key step forward in the transition of DNA storage from static archiving to dynamic interaction. By incorporating hierarchical organization and modular design, DynaByte enables flexible digital file management at the molecular level, laying a technological foundation for future rewritable and controllable molecular information systems.

The core design of DynaByte centers around a modular architecture that organizes information into structured groups of DynaBytes. In the current implementation, we designed six distinct types of DynaBytes—address-bytes, content-bytes, ECC-bytes, functional-bytes, flanking-bytes and control-bytes—which may be assembled like building blocks to enable scalable and flexible data structures (Fig. S7a). This modularity allows for easy customization of complexity and capacity. For example, expanding the address-byte pools from 100 to 1,000 entries per dimension increases the address space from ∼MB to ∼GB scale (100³ → 1,000³). Meanwhile, if we introduce another dimension of the address, then the space could reach 1,000^4^ (Terabyte, TB). Similarly, multi-level chaining of content-bytes (e.g., from one to four levels) quadruples the data payload per unit. Adding new types of functional-bytes such as version-bytes and filename-bytes enables version control and file-level metadata encoding. Importantly, the ligation-based assembly strategy used in DynaBytes can be joined in series — with literature demonstrating the successful ligation of dozens of DNA fragments [29], allowing the system to accommodate larger data blocks or more complex strucures without redesigning individual components.

Unlike most existing DNA data storage systems that are inherently read-only [30, 31], DynaByte Pro supports efficient in-DNA logical rewriting. While biochemical editing methods such as gBlock synthesis, OE-PCR, or CRISPR-Cas systems have been proposed for physical rewrites [32, 33], they face challenges including complex probe design, high cost and potential off-target effects. To address this, DynaByte Pro adopts an append-only strategy, whereby edits—additions, deletions or modifications—are encoded as incremental differences rather than overwriting the original sequence. This approach is facilitated by version-bytes, enabling edit traceability and efficient logical rewriting without compromising data integrity.

Furthermore, control-bytes, including plasmid carriers and directory tags, allow for the construction of a hierarchical file directory within DNA. Compared to magnetic bead–based or large primer-pool–based random access strategies [34–36], DynaByte Pro supports multi-level addressability—from directory to file to block—through logical address structures embedded in the DynaByte sequences. The plasmid-based architecture also offers enzymatic accessibility for future *in vitro* editing and is compatible with diverse “DynaByte” libraries, enhancing both adaptability and scalability. The accompanying modular codec pipeline also follows the modular design principle, supporting flexible encoding, decoding and logical operations (Fig. S7b).

To ensure data robustness, DynaByte employs a lightweight one-dimensional redundancy scheme (Outer Code) combined with fuzzy decoding at the module level, reducing the required sequencing depth to 100× or lower. Compared to traditional two-dimensional codes [9, 37], this approach offers lower computational cost while maintaining reliable data recovery. The modular “DNA fragment–bitstream” design provides inherent fault tolerance and synthesis compatibility, avoiding polymerization issues and reducing sensitivity to base-level errors such as substitutions and indels. This makes DynaByte particularly well-suited to real-time, high-noise platforms like ONT, where high-accuracy basecalling enables decoding performance comparable to NGS. The strong compatibility with ONT also lays the groundwork for future real-time reading and streaming-based decoding.

On the writing end, we estimate that the current prototype can achieve an average speed of ∼5.37 bytes per second, a rate expected to increase with higher-throughput synthesis and automation. Moreover, as all DynaBytes are ligated simultaneously to form a single storage unit, allowing sample processing to proceed without waiting for sequental ligaion reactions and avoiding additional time penalties. Furthermore, DynaBytes are pre-fabricated and their assembly relies on a single enzyme, which is expected to significantly reduce both laboratory workloads and production costs as the method moves toward industrial applications.

## Conclusion

Overall, this study presents a flexible molecular information system prototype, demonstrating that DNA can serve not only as a dense storage medium but also as a substrate for file system–like logic. Notably, the high flexibility achieved by DynaByte come at the expense of reduced storage density. This trade-off prompts us to envision a hybrid DNA storage architecture that combines the modularity and rewritability of DynaByte with the compactness of synthetic approaches, aiming to balance flexibility and storage efficiency. Such a system could leverage high-density methods for cold data archiving, while using DynaByte for rapid writes, versioning and index-driven access—addressing the need for responsiveness and diversity in future molecular information systems. Key challenges remain in standardizing module interfaces, developing a unified decoding framework and controlling assembly costs. Nevertheless, this direction may drive DNA storage beyond static archiving toward intelligent, functionally diverse molecular infrastructures.

## Materials and Methods

### Design of DynaByte structure and codebook

DynaBytes are short double-stranded DNA fragments with sticky or blunt linkers at both ends to enable directional ligation. Each fragment follows the structure Ver–X–Y–Z–Content–R, where the internal X–Y–Z–Content elements form the core storage unit. Ver and R serve as functional and flanking DynaBytes, respectively, enabling metadata labeling and unit delimitation. A comprehensive codebook was developed to map byte values, addresses, and metadata to DNA sequences. Content DynaBytes represented all 256 byte values using 20 bp sequences optimized for GC balance, absence of homopolymers or secondary structures, lack of restriction sites, and a minimum Levenshtein distance of 7. Functional DynaBytes (e.g., versioning, file labels) were drawn from the same pool and mapped to semantic tags. Address DynaBytes were constructed from triplets (X, Y, Z) using orthogonal 30 bp sequences, yielding an address space of up to one million locations. Linkers of 4–5 bp ensured exclusive ligation compatibility.

### Encoding scheme of the DynaByte system

Files were converted into binary strings and segmented into bytes. Groups of bytes were appended with Reed–Solomon error-correcting codes (ECC). Each byte was linearly indexed and mapped into a three-dimensional address space via a (X, Y, Z) conversion, supporting scalable and parallel organization. Indexed blocks were translated into DynaByte sequences through a codebook-derived hash table.

### Error correction strategy

An outer Reed–Solomon coding scheme (the DynaByte Outer Code) was applied to logical content groups. In silico simulations guided optimization of group size and redundancy under different dropout rates. A configuration of k=4 content bytes with n=2 ECC bytes (50% redundancy) was selected, providing tolerance to ∼20% dropout while balancing efficiency.

### DynaByte manipulation system

Functional DynaBytes were introduced for version control and file-level organization. Version DynaBytes allowed append-only editing by storing new content alongside originals and tagging incremental updates. Filename DynaBytes enabled file identification and multiplex querying. These modules facilitated scalable operations such as version tracking and selective amplification.

### DNA synthesis and assembly

All DynaByte sequences were synthesized as phosphorylated oligonucleotides, annealed into double-stranded DNA, and purified. For *in vitro* storage, DynaBytes were ligated into linearized plasmid backbones using T4 ligase in miniaturized reactions dispensed by automated liquid handling. For *in vivo* storage, ligation products were transformed into *E. coli* and propagated. A miniaturized paper-based assembly platform was also developed to reduce reaction volumes to 1–5 μL, supporting cost-efficient high-throughput ligation.

### Sequencing and retrieval

DynaByte pools were amplified and sequenced using Illumina (PE150) and Oxford Nanopore platforms. Data access employed multiplex and nested PCR, where version- and filename-DynaBytes enabled selective and hierarchical retrieval at the file or version level.

### Data processing and decoding

Sequencing reads were merged, trimmed, and aligned against the reference codebook. Each contig was segmented into address and content fragments, mapped to codebook sequences, and organized into the 3D address space. ECC-based correction was then applied. A fuzzy-matching algorithm using Levenshtein distance was implemented to tolerate sequencing noise, particularly in nanopore reads.

### Robustness evaluation

Random down-sampling simulations were performed at varying depths (5×–200×) to assess decoding reliability. Recovery rates and file completeness were quantified across 10 replicates per depth.

See details in Supplementary data.

## Acknowledgment

We would like to thank the Institutional Center for Shared Technologies and Facilities at the Wuhan Institute of Virology, CAS, Zhixian Qiao and Xiaocui Chai at the Analysis and Testing Center of the Institute of Hydrobiology, CAS, for their assistance with next-generation sequencing, and Dr. Yiqun Chen of Shanghai Dynegene Technologies Co., Ltd. for the help of DNA synthesis. Special thanks to Drs. Xiaobo Ren and Xin Chu of CAS for the efforts to push forward DNA data storage in CAS.

## Author contributions

Conceptualization: D.L., F.C., D.B., H.Y., Q.L., L.L., B.D.

Methodology: L.J., Y.S. (Yue Shi), J.Y., Y.Z., M.Z.

Investigation: L.J., Y.S. (Yue Shi), J.Y.,S.L., W.Y., Y.Z., M.Z., X.W., H.L., W.L.

Visualization: L.J., Y.S. (Yue Shi), J.Y.

Funding acquisition: D.L., Y.Z.

Project administration: D.L., L.J.

Supervision: D.L., Y.S. (Yongyong Shi)

Writing – original draft: L.J., Y.S. (Yue Shi), J.Y., S.L.,W.Y.

Writing – review & editing: D.L., Y.S. (Yongyong Shi)

## Supplementary data

Supplementary data includes seven figures and nine tables.

## Materials & Correspondence

Correspondence and requests for materials should be addressed to.

## Data availability

Raw sequencing data is deposited at https://doi.org/10.5281/zenodo.15523009 [38]. Digital files stored using DynaByte, their encoded DNA sequences, and recovered outputs can be found at https://doi.org/10.5281/zenodo.15532239 [39]. All other data are available on request.

## Code availability

The code can be found at https://doi.org/10.5281/zenodo.15550942 [40].

