## Supplemental Figures for "DNA Data Storage Architecture via Ligation of Dynamic DNA Bytes"

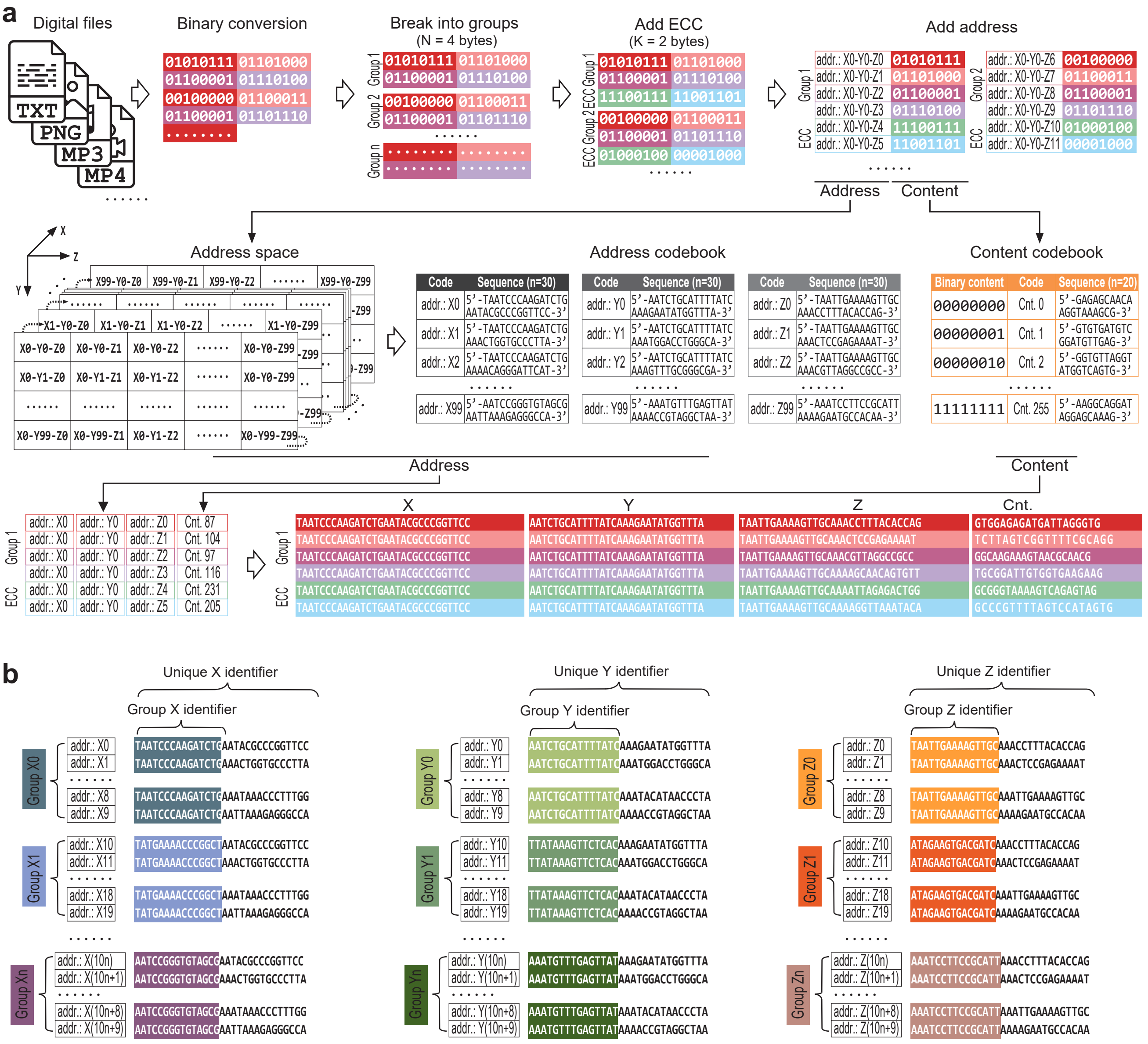


**Figure S1. Schematic of DynaByte Encoding.**

**a**, DynaByte encoding workflow, illustrating the conversion of digital information into DynaByte format and corresponding DNA sequences. **b**, Design of composite address index sequences. Each address level contains 100 unique sequences, divided into 10 groups with 10 sequences per group. Each 30 bp address is composed of a 15 bp prefix and a 15 bp suffix. Sequences within the same group share the same prefix, ensuring intra-group consistency for block-wise organization. The suffixes are unique within each group and reused across groups. This modular design uses only 20 distinct 15 bp sequences (10 prefixes + 10 suffixes) to generate 100 distinct addresses, balancing intra-group similarity and inter-sequence specificity for efficient block partitioning and sequence retrieval.

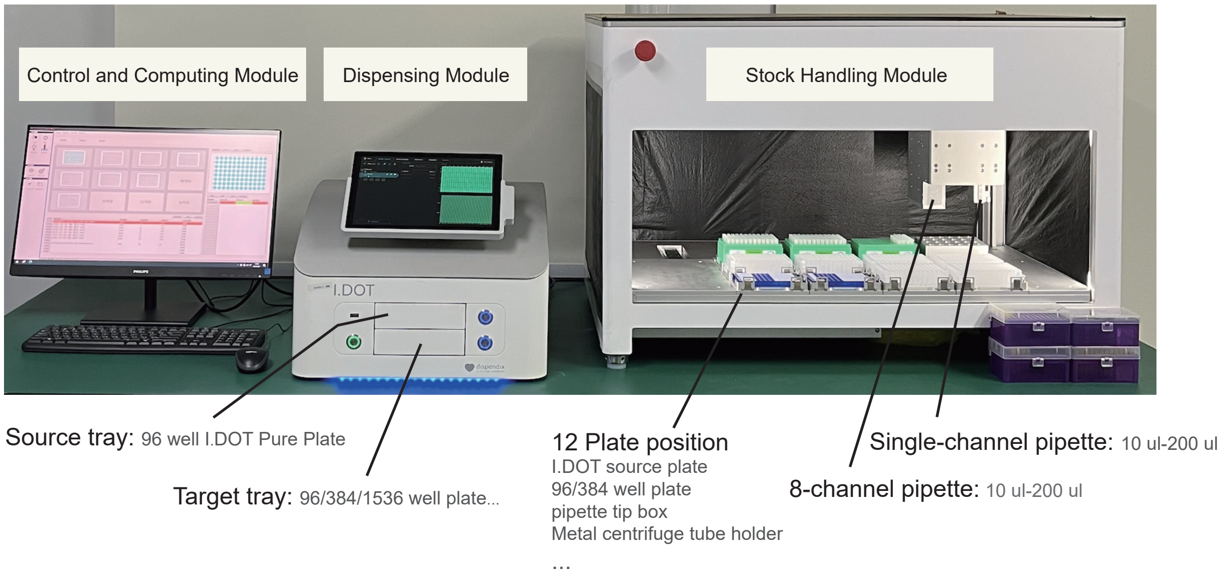


**Figure S2. Setup of the DynaByte Storage System.**

Photograph of the automated DynaByte storage system, consisting of three modules: control and computing module, stock handling module, and dispensing module. The dispensing module utilizes the I.DOT non-contact dispenser (Dispendix GmbH) for high-throughput, precise liquid handling.

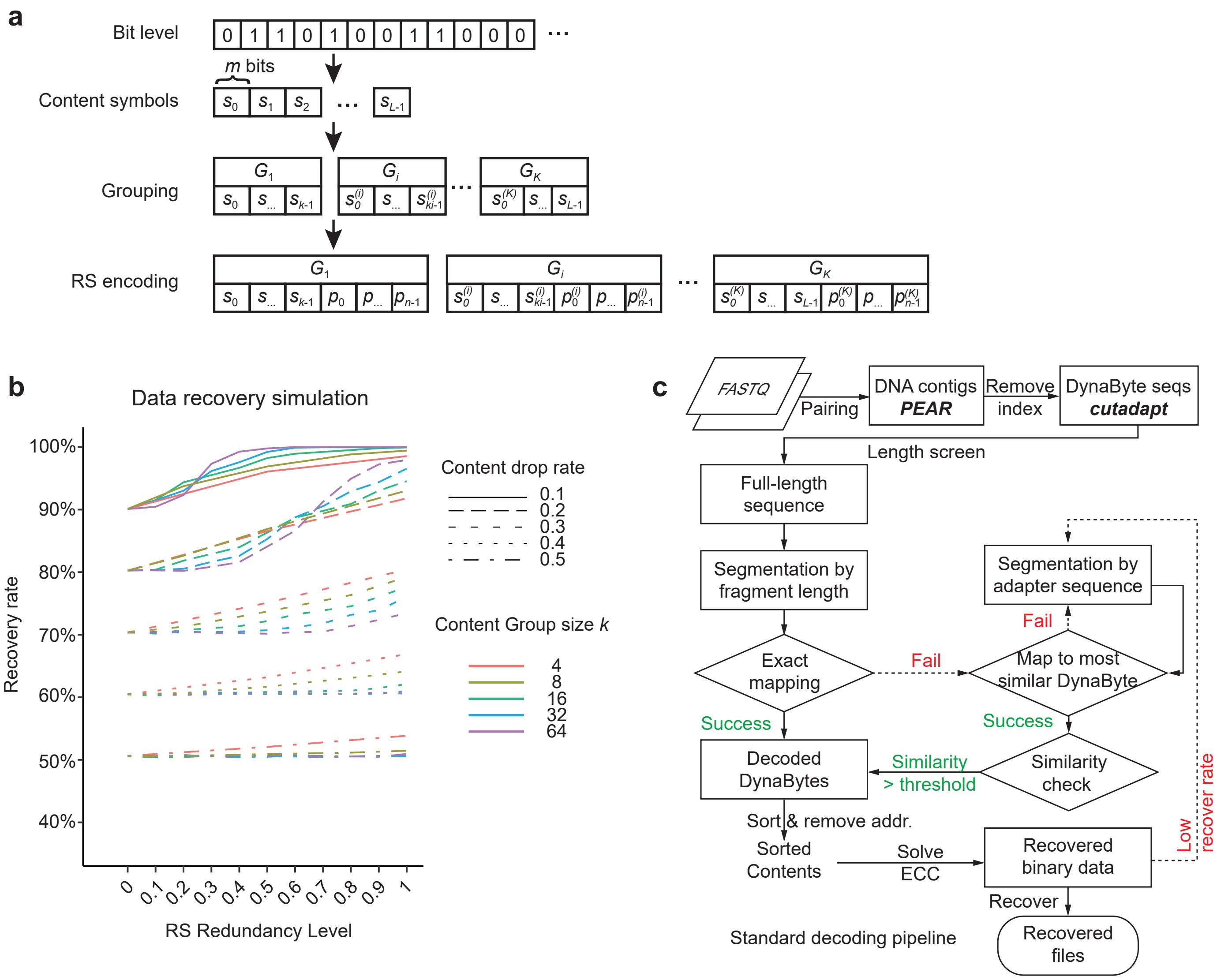


**Figure S3. ECC-byte Encoding Structure, Simulation of Recovery Performance, and Bioinformatics Pipeline for DynaByte Decoding.**

**a**, Schematic of parameter structure for ECC-byte encoding. The original bitstream is divided into fixed-length content elements (symbols) and organized into groups for Reed–Solomon (RS) error correction. Key parameters include: element bit length ($m$), total number of symbols ($L$), number of groups ($K$), group size ($k$), number of ECC symbols per group ($n$). $G_i$ denote the i-th RS encoding group. Each group consists of: $s_j$: the $j$-th data symbol (content element) in group $G_i$, where j ∈ {1, ..., k}; $p_j$: the $j$-th ECC symbol generated by RS encoding for group $G_i$, where j ∈ {1, ..., n}. **b**, Simulation of data recovery performance under different redundancy granularities. Line styles indicate content loss rates, while line colors represent RS group sizes $k$ (e.g., 4, 8, 16, 32, 64 elements per group). The x-axis denotes RS redundancy level, and the y-axis shows recovery rate. **c**, Bioinformatics pipeline for DynaByte decoding from raw sequencing data.

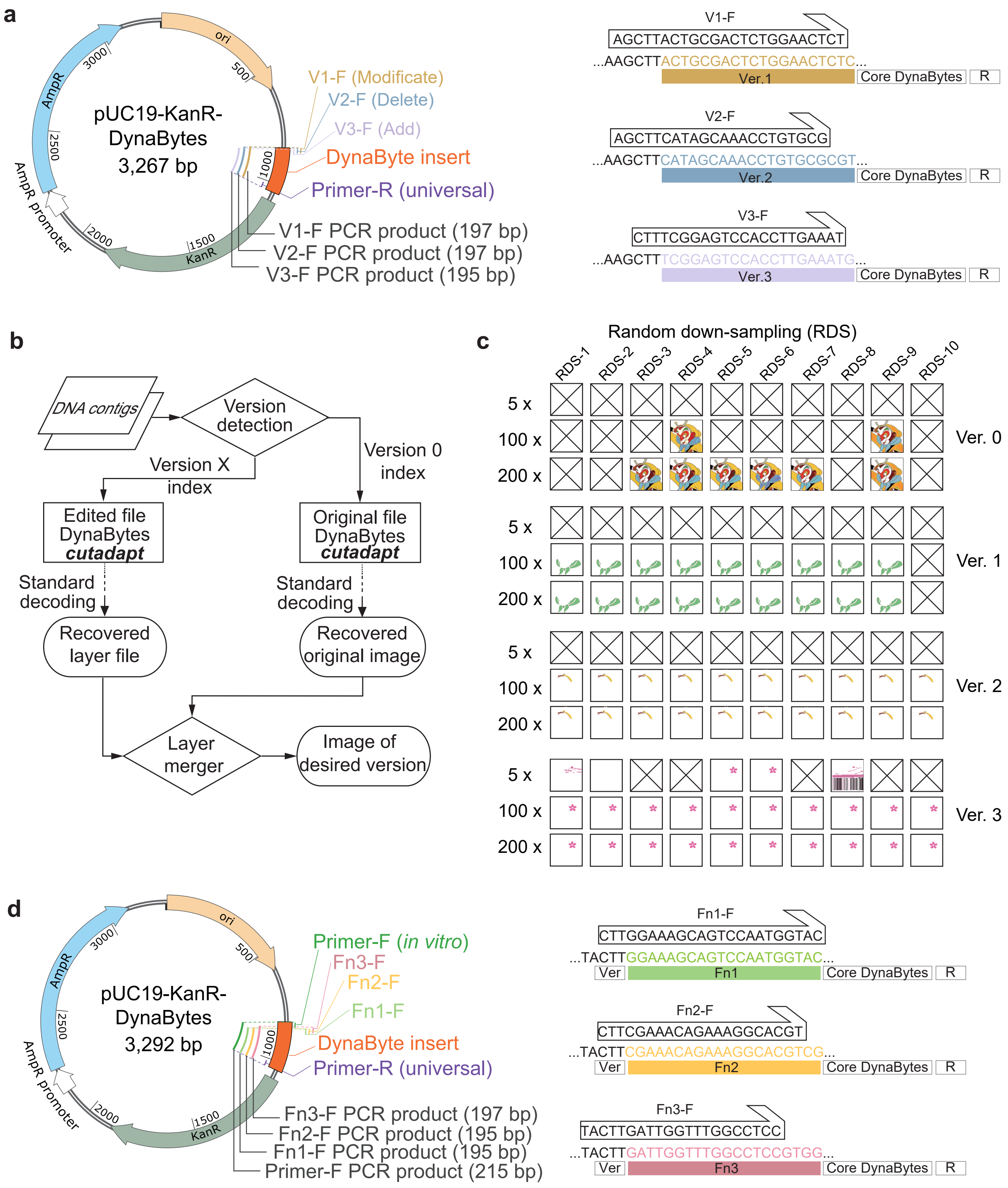


**Figure S4. Experimental and bioinformatics workflows for key-based file editing and selector-based file retrieval in DynaByte.**

**a**, Key sets and expected amplicon sizes used in file editing experiments, annotated on a plasmid map. Highlighting indicates the binding position and exact sequence of the upstream key used for targeted amplification. **b**, Bioinformatics pipeline for data decoding following file editing operations. **c**, Recovery performance of the original and edited layers—scarf color modification, hairband deletion, and flower addition—under varying sequencing depths, based on *in silico* down-sampling of NGS data. **d**, Selector sets and expected product sizes used in file search experiments, shown on a plasmid map. Upstream selector positions and sequences are specifically highlighted.

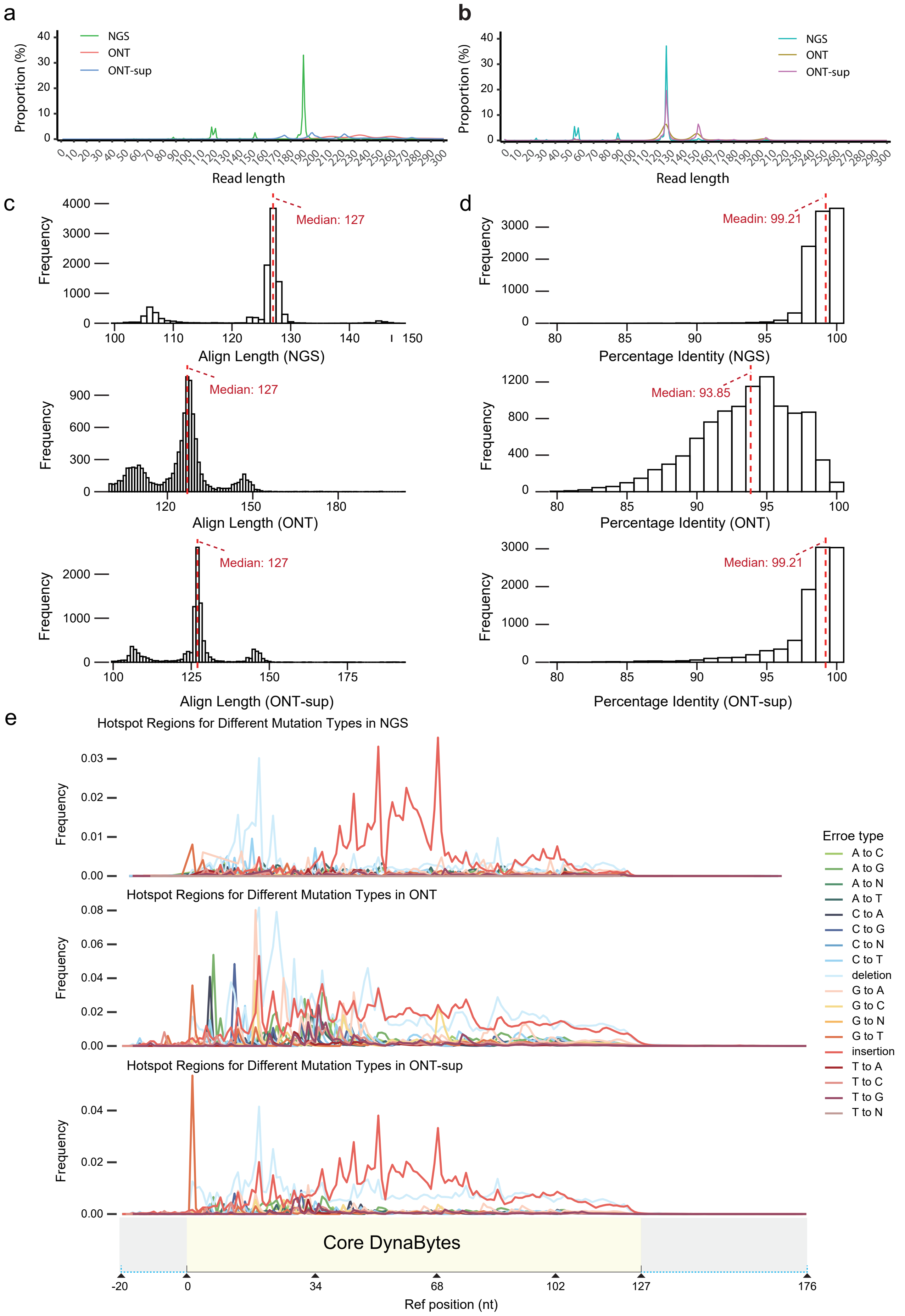


**Figure S5. Read Length, Alignment, and Error Profile Analysis for Different Reading Methods.** **a**, Distribution of read lengths after trimming version-byte across different sequencing platforms. **b**, Distribution of read lengths after further trimming the terminal R DynaBytes (flanking-byte). **c**, Alignment length distribution of sequencing reads against reference sequences using BLASTn. **d**, Identity distribution of BLASTn-aligned sequencing reads compared to reference sequences. **e**, Distribution of base-level error types across the read positions for different sequencing methods. Line colors denote error categories (e.g., mismatches, insertions, deletions).

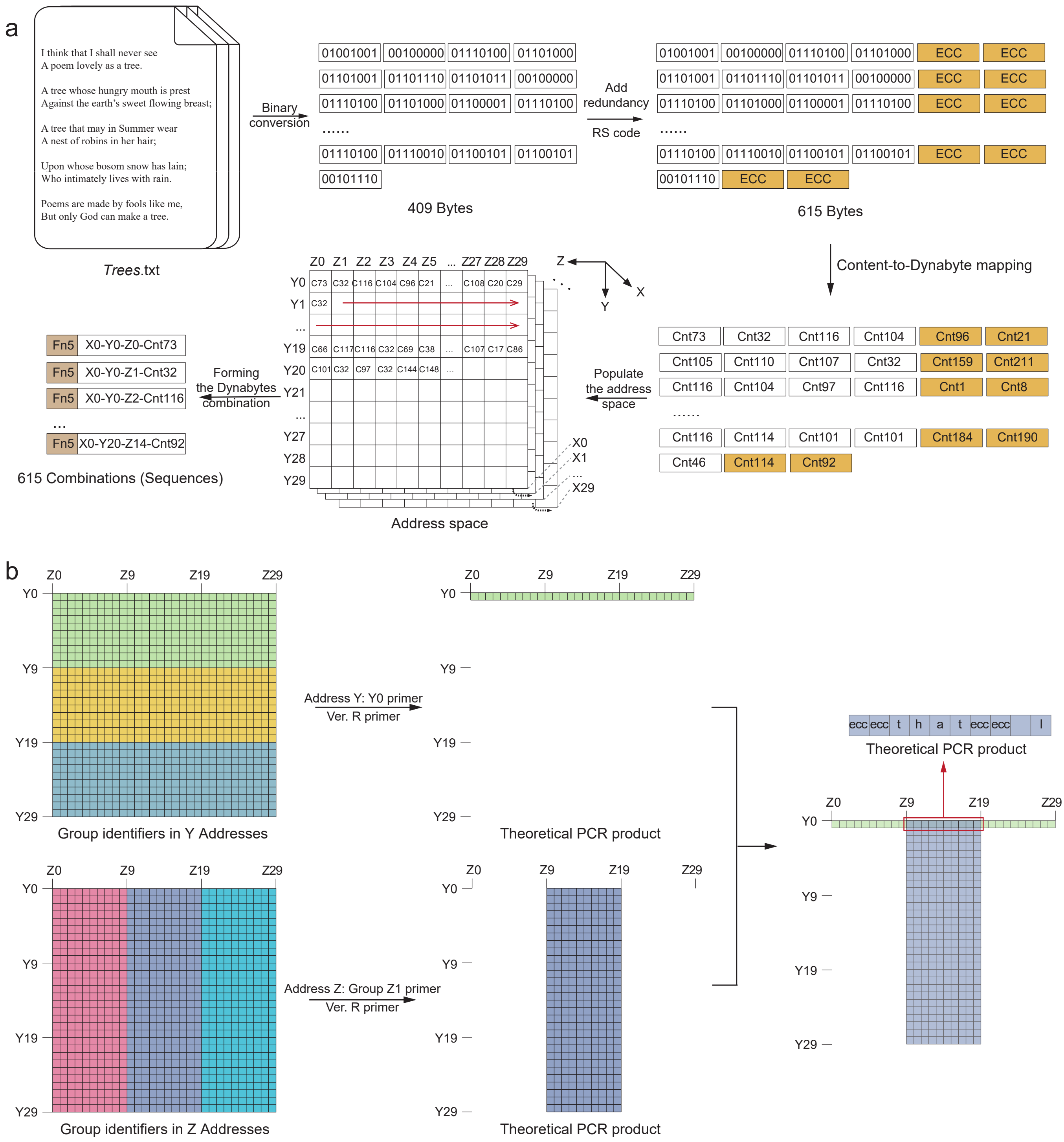


**Figure S6. Hierarchical encoding and block-wise retrieval in DynaByte Pro storage.**
**a**, Encoding process of *Trees*.txt into DynaBytes, demonstrating the incorporation of hierarchical address information. **b**, Stepwise retrieval of specific content blocks using nested PCR, targeting address Y = Y0 and Z = Z9 to Z19.

a Glossary of DynaByte bricks

| Full term | Abbreviation | Description |
|---|---|---|
| Content DynaByte | Content-byte | Encodes the actual data content. |
| Address DynaByte | Address-byte | Specifies the logical or physical storage address within the data system. |
| Error Correction Code DynaByte | ECC-byte | Introduced to support error correction. |
| Functional DynaByte | Functional-byte | Refers to DynaBytes that provide essential functions, such as version control (version-byte) or file identification (fname-byte), helping to organize and manage the stored data. |
| Flanking DynaByte | Flanking-byte | Marks the begining or end of a unit, helping to define and sta-bilize the data during storage and retrieval. |
| Control DynaByte | Control-byte | Refers to DynaBytes used as external control elements, such as primers and plasmid carrier. |

b

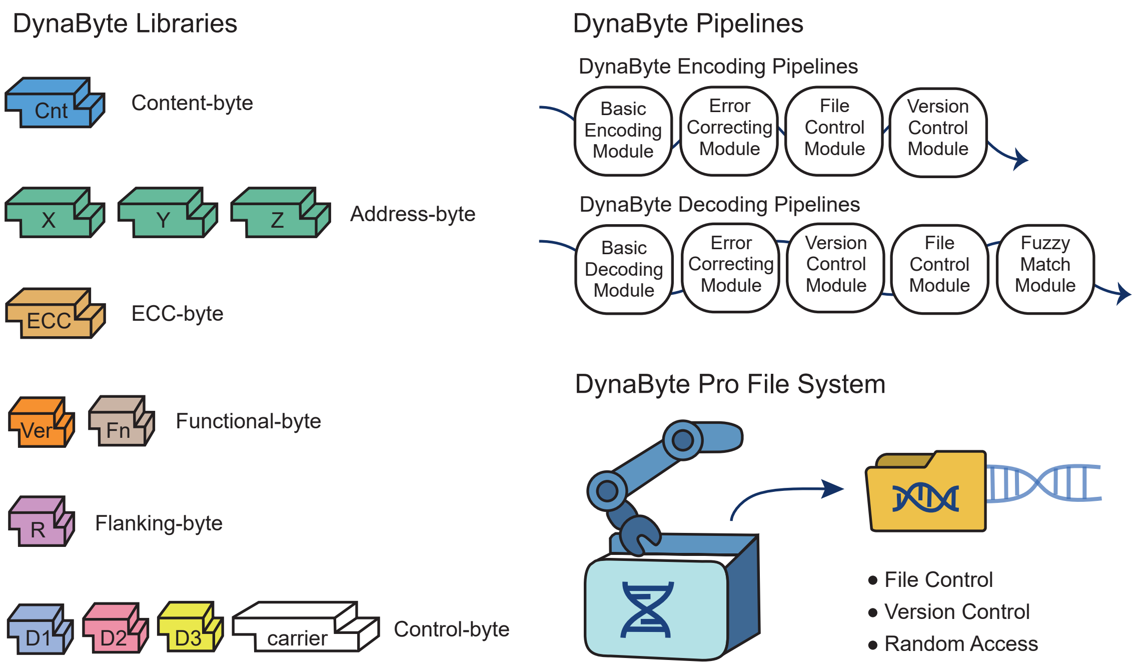


**Figure S7. Modular design of the DynaByte storage system.**
**a**, Diverse types of DynaBytes were designed to support different storage functions, including basic encoding, file-level control, versioning, fuzzy redundancy, and logical addressing. **b**, A modular DynaByte libraries and a suite of customizable DynaByte pipelines together form the flexible foundation of the DynaByte pro file system. These modules are plug-and-play and can be freely assembled to meet application-specific requirements such as file segmentation, version control, random access, and error-tolerant decoding.

### Supplemental Tables

**Table S1. The Reference DynaByte Codebook.**

This codebook defines the full sequence space and structure of DynaBytes designed for data encoding, error correction, and functional operations. All functional Sub-Codebooks used in this study were derived from this reference set.

**Table S2. Sub-Codebook A used for encoding the TXT file "What can I hold you with".**

DynaByte sequences and mapped values are used for text storage.

**Table S3. Sub-Codebook B used for encoding the PNG file "Wukong".**

DynaByte sequences and mapped values are used for picture storage.

**Table S4. Decoding Outcomes from Random Subsampling at Different Sequencing Depths.**

Statistical results from 10 independent random subsampling trials under various sequencing depths, showing decoding success rates and sequence recovery performance.

**Table S5. Sub-Codebook C used for file labeling (Fname 1-3).**

DynaByte sequences used for filename labeling, including associated metadata.

**Table S6. Sub-Codebook D used in DynaByte Pro (Fname 4-13).**

List of DynaByte sequences assigned to hierarchical directory tags and filename identifiers in the DynaByte Pro storage system, including corresponding sequences and their mapped values.

**Table S7. Average Decoding Outcomes from 10 Random Subsampling trials per file under different sequencing depths.**

For each stored file in this study, decoding was performed 10 times at each sequencing depth. The table reports the mean recovery rate for each file at each depth.

**Table S8. Decoding statistics for each digital file stored in the DynaByte system.**

This table summarizes key metrics during the decoding process of all stored files, including the number of reads retained at each processing step, decoding success rates under different matching strategies, and final recovery rates.

**Table S9. List of Primers Used in This Study.**

This file provides all primer sequences (keys or selectors) used for amplification, editing, and retrieval of DynaByte-encoded data, along with relevant annotations.
